# *Drosophila* dormancy independently regulates huddling and aggregation site selection

**DOI:** 10.64898/2026.11.20.768027

**Authors:** Seth R. Odell, Ritika Joshi, Jilian L. Morejon, Joaquin Diaz-Rossignoli, Nilay Yapici Yapici, Matthew R. Meiselman

## Abstract

Dormancy is a reversible physiological state characterized by attenuated metabolism, reproduction, and feeding, yet the behavioral adaptations accompanying this state remain poorly understood. Here, we show that cold-induced dormancy in *Drosophila melanogaster* promotes the formation of large huddles (≥5 flies), which are rarely observed under warm conditions. Dormancy simultaneously shifts these huddles away from food, though the increased propensity to huddle persists even in the absence of food, revealing separable changes in huddling propensity and aggregation site selection. We further show that dormancy resets appetite, suppressing feeding despite energetic need. Neither starvation, unrestricted feeding, nor refeeding after starvation under warm conditions reproduced the dormancy-associated pattern of aggregation, suggesting that appetite suppression during dormancy extends beyond the range of normal warm hunger and satiety states. Increasing appetitive drive during dormancy nevertheless redirected aggregation toward food. A screen of appetite-regulating neuropeptide pathways identified Neuropeptide F (NPF) signaling as a regulator of both feeding and huddling during dormancy. Broad NPF activation increased feeding and suppressed huddling, whereas activation of an NPF-expressing fan-shaped body sub-population suppressed huddling without increasing feeding. Together, these findings establish huddling as a behavioral hallmark of *Drosophila* dormancy and demonstrate that the propensity to huddle can be experimentally dissociated from appetite-dependent aggregation site selection.

## Introduction

Animals rely on both physiology and behavior to maintain function in the face of environmental stress. Desert mammals conserve water through a combination of renal concentration and nocturnal foraging, while reptiles bask to elevate body temperature after cold nights (1, 2). In insects, cold exposure often reduces metabolic output, but many species also seek shelter, aggregate, or change foraging behavior (3). These physiological and behavioral adaptations act in concert to optimize energy use, reduce risk, and increase fitness under challenging conditions. Understanding how they are coordinated requires examining not only physiological changes but also the behaviors expressed during these states.

In response to cold conditions (<14 °C), *Drosophila melanogaster* attenuates aging, metabolism, and egg production in a state referred to as either dormancy or diapause (4). Unlike the hormonally committed developmental diapause observed in many insects, *Drosophila* dormancy is facultative and rapidly reversible (5), raising questions about whether it represents a coordinated physiological state or a generalized response to cold stress (6).

Physiological states are characterized by sustained, regulated conditions with distinct traits and regulatory mechanisms (7). *Drosophila* dormancy exhibits several such features, including persistence under defined environmental conditions and broad physiological reprogramming (4). However, its characteristic phenotypes, including reproductive arrest, appetite suppression, altered energy stores, and increased sleep, are highly interconnected through metabolic and endocrine regulation (5, 8). Consequently, these measurements alone provide limited means of distinguishing coordinated state regulation from the direct consequences of cold-induced metabolic suppression.

Behavior provides an independent means of addressing this problem. Physiological states can reorganize how animals process sensory information and select among competing actions, producing distinct behavioral signatures that reflect active neural regulation rather than changes in metabolic rate alone (9, 10). Despite extensive characterization of the physiological changes accompanying *Drosophila* dormancy, comparatively little is known about its behavioral organization. Here, we identify huddling (aggregation of 5 or more flies) as a behavioral hallmark of dormancy and show that dormant flies exhibit two separable changes in social organization: an increased propensity to form large huddles and a redistribution of those huddles away from food. Increased huddling persists in the absence of food, demonstrating that it is not a consequence of altered feeding behavior alone. We further show that dormancy recalibrates feeding homeostasis, producing an appetite state not reproduced by normal warm hunger and satiety, while appetitive drive nevertheless determines where flies aggregate. Manipulation of appetite-regulating neuropeptide pathways identified NPF as a potent regulator of feeding and huddling during dormancy. Broad NPF-neuron activation increased feeding and suppressed huddling, whereas activation of a subset of NPF neurons projecting to the fan-shaped body (NPF^FSB^) suppressed huddling without increasing feeding. Together, these findings establish huddling as a regulated behavioral component of dormancy and demonstrate that huddling propensity and appetite-dependent aggregation site selection are experimentally separable.

## Results

### Dormancy alters social behavior independently of locomotor suppression

To identify behavioral signatures of dormancy, we continuously tracked freely behaving groups of *Drosophila melanogaster* maintained under warm (25°C) or dormancy-inducing cold (12°C) conditions using a custom overhead imaging system (LOOC) over a 12:12 light:dark cycle (11). Consistent with previous measurements obtained using the Drosophila Activity Monitor (DAM) system (8, 12), flies housed at 12°C exhibited a profound reduction in circadian locomotor activity, including attenuation of the characteristic morning and evening activity peaks, and a greater than 75% reduction in walking velocity as measured by multi-animal tracking (T-Rex) (Fig. 1A-B) (13). To determine whether dormancy also altered social behavior, we used a custom position-tracking software paired with T-Rex data (LOOCAPT) to calculate each fly’s average distance to all cohort members as a continuous measure of grouping behavior (Fig. 1C-D). In warm conditions, inter-fly distance fluctuated with locomotor activity over the circadian cycle, whereas dormant flies maintained relatively constant group spacing throughout the day regardless of activity (Fig. 1E, Fig. S1A-B). Because reduced activity was associated with decreased inter-fly distance, and sleep is altered during dormancy (8), we asked whether manipulating sleep would also affect grouping. Neither caffeine treatment during dormancy nor sleep deprivation under warm conditions significantly altered grouping distance, suggesting that reduced activity and increased sleep alone do not account for the altered spatial organization (Fig. S1C-F).

**Fig. 1.**
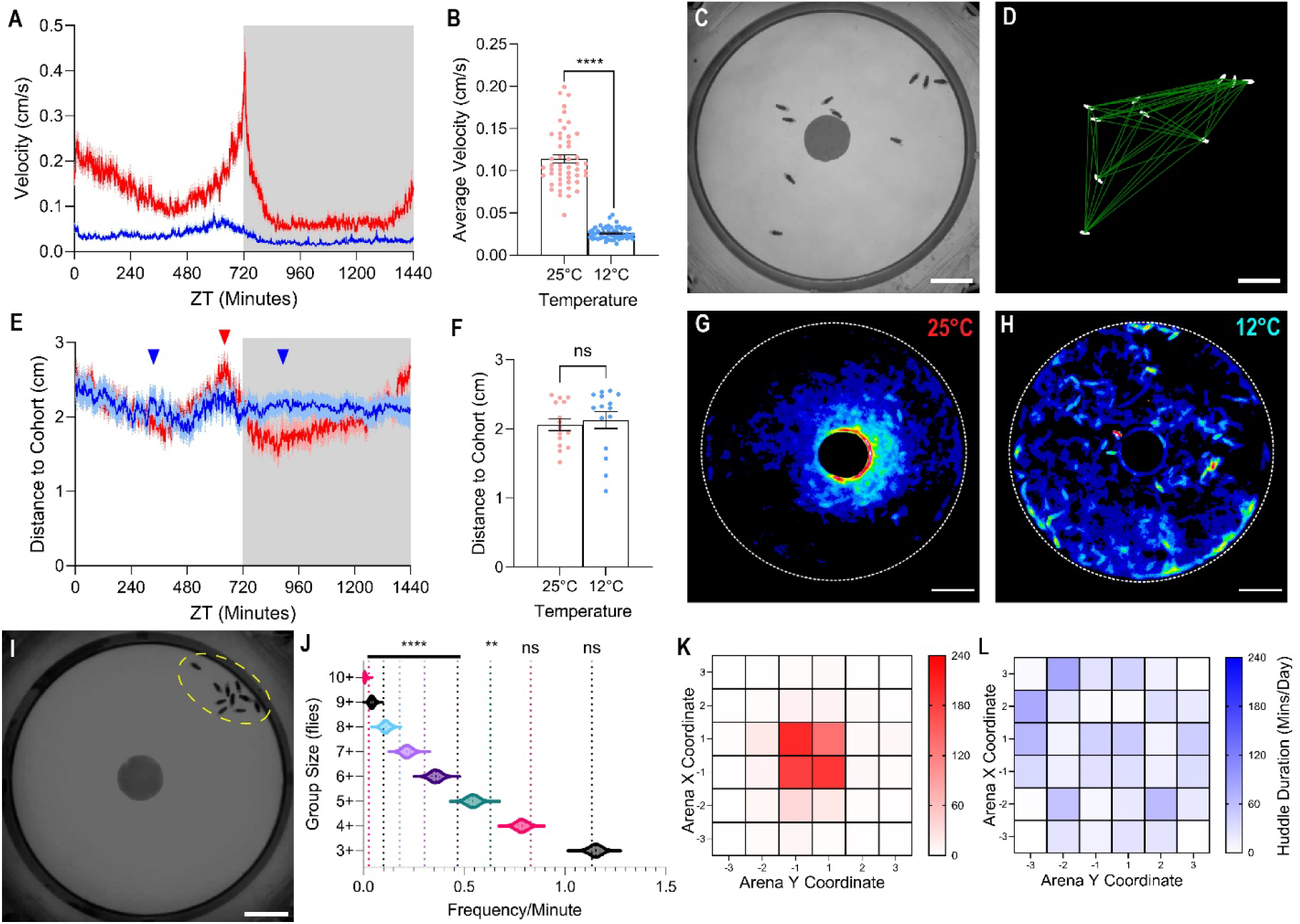
Dormancy suppresses locomotor activity while reorganizing spatial occupancy. **(A)** Mean locomotor velocity of flies maintained at 25°C (red) or dormancy-inducing 12°C (blue) over a 24-hr 12:12 light:dark cycle. Gray shading indicates the dark phase; dashed lines indicate ± SEM (*n* = 50-51 flies). **(B)** Mean locomotor velocity over the 24-hr recording period at 25°C or 12°C. Each point represents one fly (*n* = 50-51 flies); bars indicate mean ± SEM. Student’s *t* test,****, *P* < 0.0001. **(C and D)** Representative frame **(C)** of the recording arena showing the central food source and fly positions. **(D)** Example multi-animal position tracking used to generate distance-to-cohort measurements, with flies shown in white and inter-fly distances shown as green lines. Scale bars, 1 cm. **(E)** Mean distance to cohort at 25°C (red) or 12°C (blue) over a 24-hr 12:12 light:dark cycle. Arrowheads indicate periods of elevated distance to cohort for flies in cold (blue) or warm (red). Gray shading indicates the dark phase; lines indicate mean and shading indicates ± SEM. **(F)** Mean distance to cohort over the 24-hr recording period. Each point represents one independent trial (*n* = 15 trials per temperature); bars indicate mean ± SEM. Student’s *t* test, *ns*. **(G and H)** Representative heat maps showing cumulative spatial occupancy over 24 hr for flies maintained at 25°C **(G)** or 12°C **(H).** Scale bars, 1 cm. **(I)** Representative frame showing a huddle (green ellipse). Scale bar, 1 cm. **(J)** Randomization analysis used to establish the operational definition of a huddle. Observed frequencies of groups containing 3-10 flies were compared with frequencies generated from randomized fly positions. Groups of five or more flies occurred at frequencies exceeding the 95th percentile of the randomized distribution and were therefore classified as huddles. Significance was determined relative to the randomized distribution, as described in Materials and Methods. **(K and L)** Spatial distribution of huddle centroids at 25°C **(K)** or 12°C **(L)** across a 6 × 6 arena grid. Each huddle was assigned to the grid square containing its centroid; color indicates the cumulative huddle duration (minutes per 24-hr recording) within each grid square.

Consequently, overall average inter-fly distance over 24 hours was similar between temperatures despite markedly different circadian and temporal dynamics (Fig. 1F).

To examine the spatial organization underlying these differences in grouping behavior, we created heat maps of fly positions in the arena in warm and cold conditions over 24 hours at warm or cold temperatures (Fig. 1G-H). We observed a dramatic shift in regional occupancy between warm and cold conditions; warm flies grouped near food, whereas dormant flies appeared to disperse more evenly through the arena. Occupancy maps also revealed “hot spots,” located in the center in warm and near edges in cold. When we reviewed videos, we found large groups of static flies in both conditions (Fig. 1I). Comparison with randomized fly positions showed that groups of five or more flies occurred more frequently than expected by chance (Fig. 1J). We therefore defined these large groups as “huddles”. Huddle centroids correlated with overall occupancy, with huddling occurring near food in warm and evenly dispersed in cold (Fig. 1K-L).

### Dormancy Independently Regulates Huddling and Aggregation Site Selection

To quantify changes in regional occupancy, we divided the arena into two zones: an annulus (donut-shaped area) extending approximately two fly body-lengths from the food edge (Feeding Zone, 3.37 cm^2^), and the area outside (Margin Zone, 29.02 cm^2^) (Fig. 2A). We compared huddle participation in each region over 24 hours, finding that Feeding Zone huddling was ∼5-fold more frequent in warm conditions (21.42 ± 3.33% SEM at 25°C versus 4.11 ± 2.35% at 12°C), whereas Margin Zone huddling was ∼8-fold more frequent in cold conditions (4.61 ± 0.92% at 25°C versus 38.73 ± 6.23% at 12°C) (Fig. 2B-E). These differences persisted during both light and dark periods. Consistent with a limited contribution of visual input, Margin Zone huddling was largely preserved in visually impaired *ort*, *ninaE* mutants (Fig. S2A-F). In contrast, *Orco* mutants exhibited reduced Margin Zone and total huddle participation during dormancy. Because olfaction also contributes to food-seeking behavior, we asked whether this reduction was accompanied by altered feeding, but detected no significant difference between *Orco* mutants and controls (Fig. S2A-G).

**Fig. 2.**
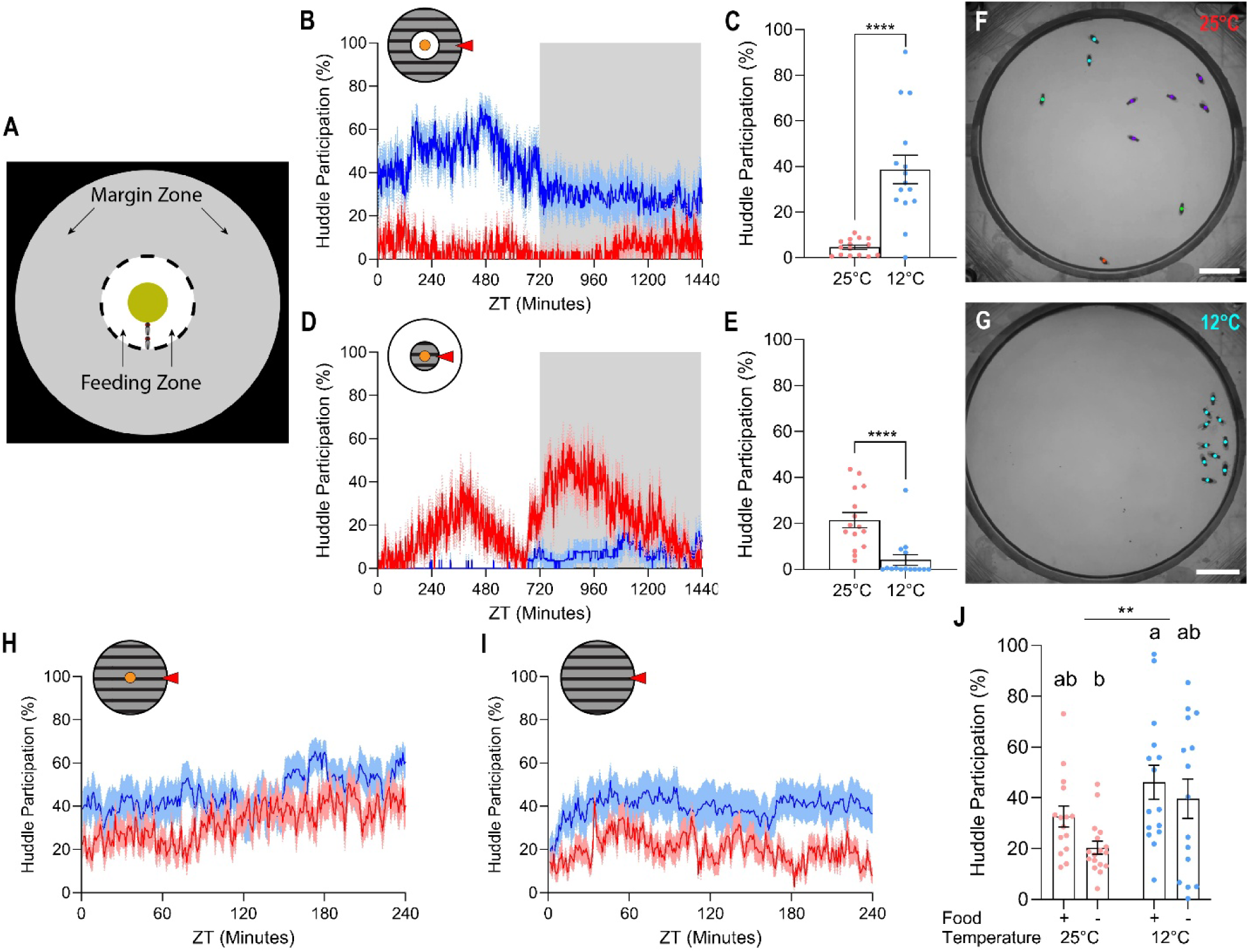
Dormancy increases huddling while shifting aggregation away from food. **(A)** Schematic of the behavioral arena. The food disc (0.5-cm radius) was positioned centrally within a 3.25-cm radius arena. The Feeding Zone was defined as the annulus around the food disk approximately two fly body lengths (1.15-cm outer radius), with the remaining area defined as the Margin Zone. **(B)** Mean participation in huddles centered in the Margin Zone (red arrow and shading in inset) over a 24-hr 12:12 light:dark cycle at 25°C (red) or dormancy-inducing 12°C (blue). Gray shading indicates the dark phase; lines indicate mean and shading indicates ± SEM (*n* = 15 trials per temperature). **(C)** Mean percentage of flies participating in Margin Zone huddles over the 24-hr recording period. Each point represents one independent trial (*n* = 15 trials per temperature); bars indicate mean ± SEM. Student’s *t* test, ****, *P* < 0.0001. **(D)** Mean participation in huddles centered in the Feeding Zone (red arrow and shading in inset) over a 24-hr 12:12 light:dark cycle at 25°C (red) or 12°C (blue). Gray shading indicates the dark phase; lines indicate mean and shading indicates ± SEM (*n* = 15 trials per temperature). **(E)** Mean percentage of flies participating in Feeding Zone huddles over the 24-hr recording period. Each point represents one independent trial (*n* = 15 trials per temperature); bars indicate mean ± SEM. Student’s *t* test, ****, *P* < 0.0001. **(F and G)** Representative frames of the recording arena showing huddles at 25°C (**F**) and 12°C (**G**). Colored dots identify flies assigned to the same group. Scale bars, 1 cm. **(H)** Mean participation in huddles anywhere in the arena during the first 240 min of the day at 25°C (red) or 12°C (blue) in arenas containing food. Lines indicate mean and shading indicates ± SEM (*n* = 15 trials per temperature). **(I)** Mean participation in huddles anywhere in the arena during the first 240 min of the day at 25°C (red) or 12°C (blue) in arenas lacking food. Lines indicate mean and shading indicates ± SEM (*n* = 15-17 trials per temperature). **(J)** Mean huddle participation during the 240-min assay in arenas with (+) or without (−) food at 25°C or 12°C. Each point represents one independent trial; bars indicate mean ± SEM. Two-way ANOVA, significant main effect of temperature (**, *P* < 0.01). Different letters indicate significant pairwise differences (Tukey’s multiple-comparisons test, *P* < 0.05); groups sharing a letter do not differ significantly.

The olfactory dependence of huddling also raised the possibility that food-associated odors contribute to aggregation, even when huddles form away from the food source. We therefore quantified total huddling independent of region in the presence or absence of food (Fig. 2F-J). Huddling was significantly more frequent in cold than in warm conditions, both in arenas containing food and those without, demonstrating that dormancy increases the propensity to huddle independently of food availability.

### Dormancy Recalibrates Feeding Homeostasis

Flies, like many animals, are thigmotactic (14–18). We reasoned that placing a small food disk (∼2% of total arena area) in the center of the arena would limit interaction with food to periods of purposeful ingestion. We therefore sought to determine if Feeding Zone huddling was a direct representation of feeding. Using SLEAP-based pose estimation (19), we trained and automated the program to estimate the proportion of flies feeding at any given time (Fig. 3A). Automated measurements were validated against blinded manual annotations of feeding, defined by proboscis extension and cibarial pumping (20), and did not differ significantly between methods (Fig. S3A-D). In line with a previous report of reduced consumption during dormancy (5), time spent feeding was reduced from ∼13% of a fly’s day at 25°C to ∼5% in 12°C (Fig.3B). However, in both cases feeding behavior (measured by SLEAP) only accounted for 30-40% of Feeding Zone occupation, similar at both temperatures (Fig. S3E). A consistent proportion of feeding suggested altered grouping position in dormancy was at least partially driven by reduced feeding.

One explanation for reduced feeding during dormancy is that flies compensate by consuming fewer, larger meals. To test this hypothesis, we used Expresso to quantify both the proportion of flies initiating a meal and the volume consumed per meal following 24 hours of starvation (21, 22). Relative to warm flies, fewer dormant flies initiated feeding, but meal volume among flies that did consume a meal of 300mM sucrose was unchanged (Fig. 3C). Thus, the reduced feeding observed during dormancy reflects fewer flies choosing to feed rather than a change in meal size. In contrast, flies recovered from dormancy before starvation consumed meals approximately twice as large as warm flies, consistent with compensatory hyperphagia following restoration of normal feeding drive. Because meal size scales with starvation state (22), the hyperphagia observed following recovery suggests that these flies were in a more energetically stressed state than matched warm or cold flies.

**Fig. 3.**
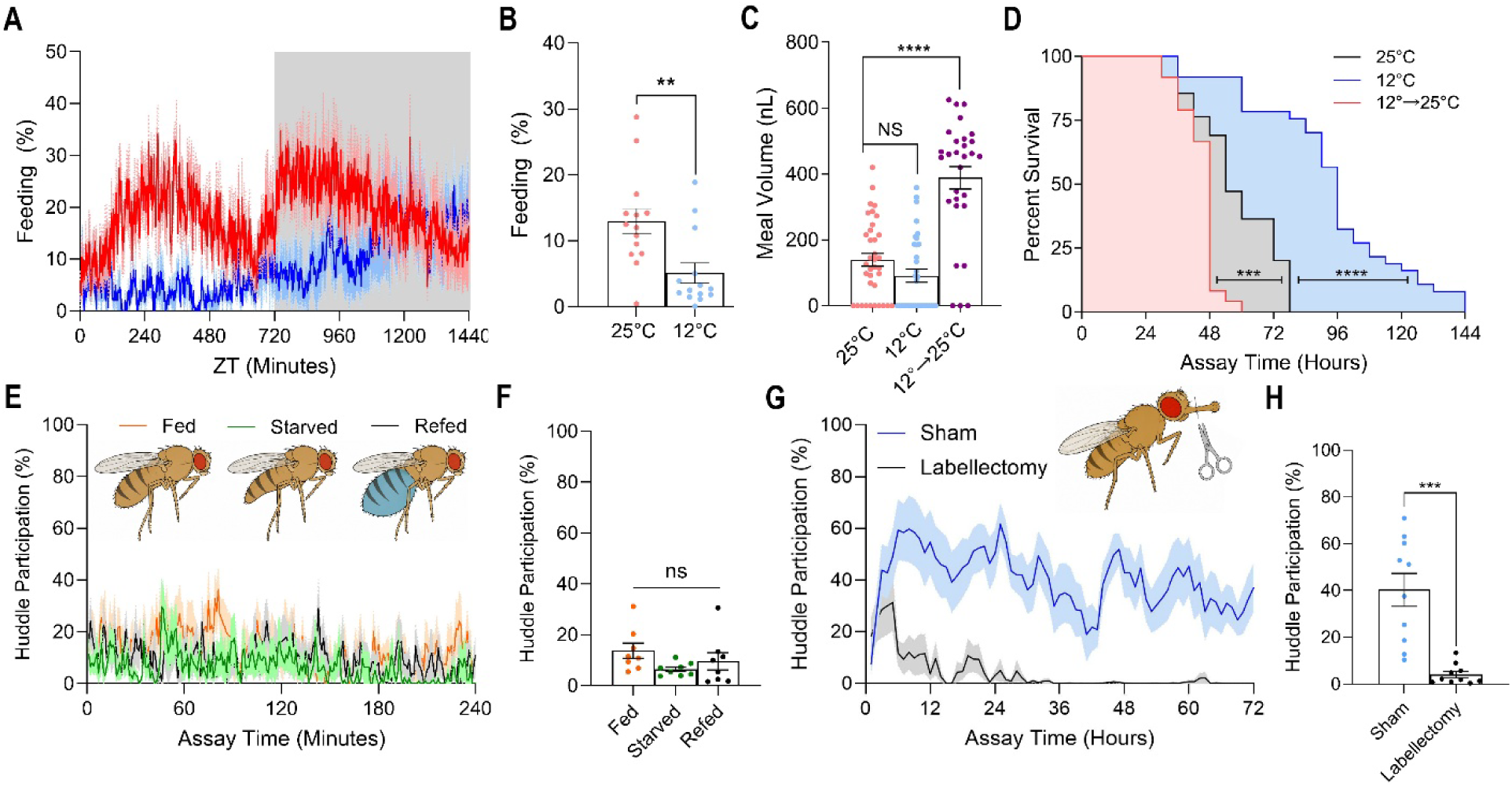
Dormancy reorganizes feeding homeostasis and appetite-dependent aggregation site selection. **(A)** Percentage of flies feeding over a 24-hr 12:12 light:dark cycle at 25°C (red) or dormancy-inducing 12°C (blue), quantified using SLEAP-based pose estimation. Gray shading indicates the dark phase; lines indicate mean and shading indicates ± SEM. **(B)** Mean percentage of time spent feeding over the 24-hr recording period at 25°C or 12°C. Each point represents one independent trial (*n* = 15 trials per temperature); bars indicate mean ± SEM. Student’s *t* test, **, *P* < 0.01. **(C)** Meal volume measured using the Expresso assay following 24 hr of food deprivation. Flies were maintained at 25°C, 12°C, or transferred from 12°C to 25°C during food deprivation (12°C→25°C). Meal volume is shown for flies that initiated feeding (*n* = 28-36 flies per condition); bars indicate mean ± SEM. Pairwise Welch’s *t* tests, *ns*; ****, *P* < 0.0001. **(D)** Survival during food deprivation for flies maintained at 25°C, 12°C, or transferred from 12°C to 25°C immediately before food deprivation (*n* = 26-55 flies per condition). Pairwise Mantel-Cox log-rank tests, ***, *P* < 0.001; ****, *P* < 0.0001. **(E and F)** Effect of acute changes in satiety on Margin Zone huddling in warm-reared flies. Flies were continuously fed (Fed), food deprived for 24 hr (Starved), or food deprived for 24 hr and allowed to feed for 10 min on 100 mM sucrose containing blue food coloring (Refed). For the Refed condition, flies exhibiting the greatest abdominal distension and visible blue coloration were selected to enrich for flies that had recently consumed a large meal, as illustrated in **E**. **(E)** Percentage of flies participating in Margin Zone huddles during the 4-hr assay. **(F)** Mean Margin Zone huddle participation over the assay. Each point represents one independent trial (*n* = 8 trials per condition); bars indicate mean ± SEM. One-way ANOVA, *ns*. **(G)** Margin Zone huddle participation following labellectomy. Dormant flies underwent labellectomy (black) or sham surgery (blue) immediately before transfer to behavioral arenas and were recorded continuously for 72 hr at 12°C. Lines indicate mean and shading indicates ± SEM. **(H)** Mean Margin Zone huddle participation over the 72-hr recording period following labellectomy or sham surgery in dormant flies. Each point represents one independent trial (*n* = 10 trials); bars indicate mean ± SEM. Student’s *t* test, ***, *P* < 0.001.

These results raised the question of whether reduced feeding during dormancy produced an energy-stressed state. We food-deprived flies at 25°C, 12°C, and in warm conditions after dormancy (12°C→25°C, recovery) (Fig. 3D). Surprisingly, flies starved while dormant survived approximately twice as long as warm-starved controls, consistent with reduced metabolic expenditure during cold exposure to match reduced overall feeding. However, flies transferred to warm conditions immediately prior to food deprivation survived significantly fewer days than warm controls despite continuous prior access to food during dormancy, suggesting that energetic demand during recovery may exceed the reserves maintained during dormancy. Together, these findings indicate that dormancy uncouples appetite from energetic state.

### Dormancy-Specific Appetite Suppression Regulates Aggregation Site Selection

Because dormant flies exhibit profound appetite suppression, we asked whether Margin Zone huddling reflects an exaggerated state of satiety. We therefore tested whether manipulating hunger and satiety in warm flies could reproduce this behavior. We measured Margin Zone huddling over four hours in warm flies under three appetitive conditions: continuously fed (Fed), food deprived for 24 hours (Starved), or food deprived for 24 hours followed by 10 minutes of refeeding on 100 mM sucrose (Refed), a concentration previously shown to maximize meal volume (22). Refed flies were selected based on abdominal distension to ensure tested individuals had consumed a large meal. Feeding Zone grouping and feeding were significantly elevated in the Starved and Refed flies compared to Fed, but Margin Zone huddling was similar in Fed, Starved, and Refed over the following four hours, and Margin Zone huddling did not approach the 40-50% participation seen in time-matched cold flies (Fig. S3F-I, Fig. 3E-F). Thus, Margin Zone huddling could not be reproduced by normal homeostatic changes in satiety, suggesting that the appetite suppression accompanying dormancy represents a distinct behavioral state.

We next asked whether increasing appetitive drive during dormancy was sufficient to suppress Margin Zone huddling. To increase feeding drive despite unrestricted food availability, we performed labellectomies, depriving flies of consumption without impacting their environment (Fig. 3G-H). Margin Zone huddling progressively declined and was rarely observed 36 hours after labellectomy, suggesting that increasing appetitive drive is sufficient to suppress dormancy-associated Margin Zone huddling.

### NPF Signaling Suppresses Dormancy-Associated Margin Zone Huddling

Recent work has implicated peptidergic neurons with known roles in both feeding and the regulation of reproductive dormancy, including DH31 (23, 24), sNPF (25, 26), AstC (27, 28), and dILPs (29, 30). Because increasing appetitive drive suppressed Margin Zone huddling, we hypothesized that peptidergic pathways regulating feeding might impact Margin Zone huddling. We therefore screened 11 neuropeptide systems implicated in appetite regulation. Five putatively anorexigenic peptides (CCH-Amide2 [CCHA2], Myoinhibitory Peptide [MIP], Leucokinin [Lk], Allatostatin-A [AstA], and Hugin) were inhibited using *UAS-TNT* (31–35). Six putatively orexigenic peptides (Insulin-Like Peptide [ILP2], short Neuropeptide F [sNPF], Allatostatin C [AstC], Neuropeptide F [NPF], Corazonin [Crz], and SIFamide [SIFa]) were activated using the cold-activated cation channel TRPM8 (25, 27, 29, 36–38).

Feeding behavior, measured by SLEAP, was significantly elevated during dormancy following activation of NPF or SIFamide neurons (*NPF-Gal4>UAS-TRPM8* and *SIFA-Gal4>UAS-TRPM8*; Fig. 4A). Three manipulations (*Lk-Gal4>UAS-TNT*, *NPF-Gal4>UAS-TRPM8* and *ILP2-Gal4>UAS-TRPM8*) significantly reduced Margin Zone huddling (Fig. 4B). Additionally, 8 of the 11 pro-appetitive manipulations increased mature egg production during dormancy relative to controls (Fig. S4A), suggesting that neuropeptide pathways associated with appetite regulation also broadly influence reproductive suppression during dormancy. Notably, whereas NPF activation both increased feeding and suppressed Margin Zone huddling, SIFa activation increased feeding without affecting huddling, while ILP2 activation and Lk inhibition reduced Margin Zone huddling without a detectable increase in feeding. Thus, feeding and huddling could be independently manipulated, demonstrating that the two behaviors are not obligatorily coupled during dormancy.

**Fig. 4.**
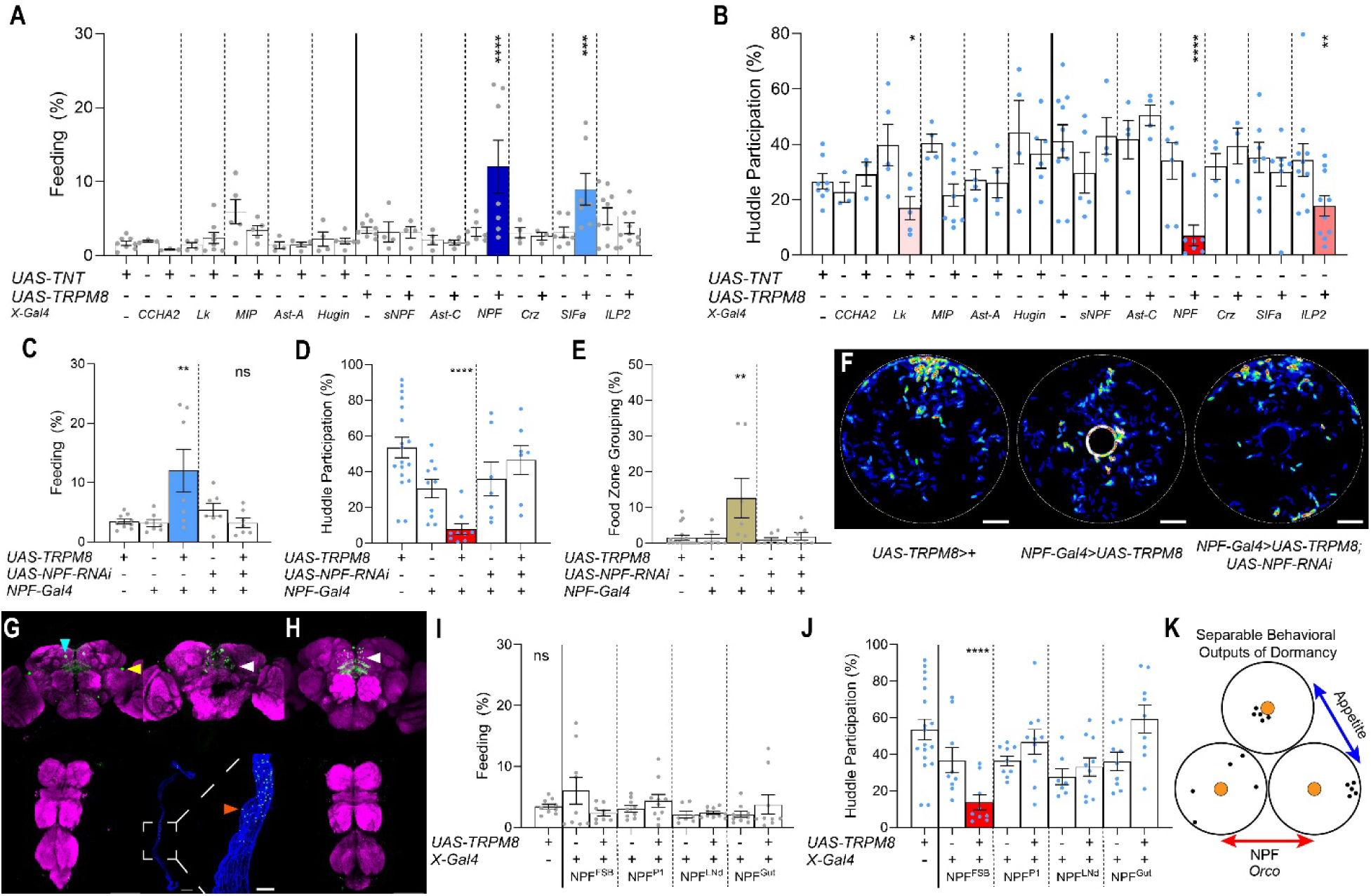
NPF signaling reveals separable regulation of feeding and huddling during dormancy. **(A and B)** Screen of appetite-regulating neuropeptide pathways during dormancy. Putatively anorexigenic pathways (CCHa2, Lk, MIP, AstA, and Hugin) were inhibited using *UAS-TNT*, whereas putatively orexigenic pathways (sNPF, AstC, NPF, Crz, SIFa, and ILP2) were activated using *UAS-TRPM8*. **(A)** Mean percentage of time spent feeding over 24 hr, quantified using SLEAP. Each point represents one independent trial (*n* = 4-10 trials per genotype); bars indicate mean ± SEM. One-way ANOVA followed by Dunnett’s multiple-comparisons test. Bar fill indicates significance relative to corresponding UAS control: white, not significant; light blue, *P* < 0.001; dark blue, *P* < 0.0001. **(B)** Mean Margin Zone huddle participation over 24 hr for the corresponding genotypes. Each point represents one independent trial (*n* = 4-10 trials per genotype); bars indicate mean ± SEM. One-way ANOVA followed by Dunnett’s multiple-comparisons test. Bar fill indicates significance relative to corresponding UAS control: white, not significant; light red, *P* < 0.05; medium red, *P* < 0.01; dark red, *P* < 0.0001. **(C-F)** Effects of simultaneous NPF activation and knockdown during dormancy. NPF neurons were activated using *UAS-TRPM8* with or without *UAS-NPF-RNAi*, alongside corresponding genetic controls. **(C)** Mean percentage of time spent feeding over 24 hr, quantified using SLEAP. Each point represents one independent trial (*n* = 7-9 trials per genotype); bars indicate mean ± SEM. **(D)** Mean Margin Zone huddle participation over 24 hr. Each point represents one independent trial (*n* = 7-18 trials per genotype); bars indicate mean ± SEM. **(E)** Mean Feeding Zone grouping over 24 hr. Each point represents one independent trial (*n* = 7-18 trials per genotype); bars indicate mean ± SEM. For **(C-E),** one-way ANOVA followed by Tukey’s multiple-comparisons test. Symbols indicate significance relative to both corresponding genetic controls, with the less significant adjusted comparison reported: *, *P* < 0.05; **, *P* < 0.01; ***, *P* < 0.001; ****, *P* < 0.0001; *ns*. **(F)** Representative 24-hr spatial occupancy heat maps of the indicated genotypes, with the central food source outlined in white. Color indicates cumulative occupancy, from low (blue) to high (red). Scale bars, 1 cm. **(G and H)** Expression patterns of *NPF-Gal4>UAS-mCD8-GFP* and *NPF^FSB^-Gal4>UAS-mCD8-GFP*. **(G)** Representative confocal images showing *NPF-Gal4* expression (green) in the adult brain viewed anteriorly (upper left) and posteriorly (upper right), ventral nerve cord (lower left), and gastrointestinal tract (lower right). Bruchpilot immunostaining (magenta) labels neuropil in the brain and VNC, whereas phalloidin staining (blue) labels F-actin in the gut. Arrowheads indicate NPF^FSB^ (white), NPF^LNd^ (cyan), and NPF^P1^ (yellow) populations. Box indicates the midgut region shown at higher magnification; arrowhead indicates NPF-positive enteroendocrine cells. Scale bars, 100 μm (brain and VNC), 500 μm (whole gut), and 100 μm (magnified midgut). **(H)** Representative confocal images of the adult brain (upper) and VNC (lower), showing *NPF^FSB^-Gal4* expression (green) and Bruchpilot immunostaining (magenta). Scale bars, 100 μm. **(I and J)** Effects of activating anatomically defined NPF subsets (NPF^FSB^, NPF^P1^, NPF^LNd^, and NPF^Gut^) using *UAS-TRPM8*. **(I)** Mean percentage of time spent feeding over 24 hr, quantified using SLEAP. Each point represents one independent trial (*n* = 7-10 trials per genotype); bars indicate mean ± SEM. **(J)** Mean Margin Zone huddle participation over 24 hr. Each point represents one independent trial (*n* = 7-18 trials per genotype); bars indicate mean ± SEM. For (I and J), one-way ANOVA followed by Dunnett’s multiple-comparisons test against the corresponding UAS controls. *ns*; ****, *P* < 0.0001. **(K)** Model illustrating two separable behavioral outputs of dormancy: appetite-dependent aggregation site selection and huddling propensity. NPF signaling influences both processes, whereas Orco-dependent olfactory signaling contributes to huddling propensity.

Broad activation of *NPF-Gal4* cells was the most effective in both categories in restoring warm-like feeding and grouping location, and moved fly position closer to the food’s edge (Fig. 4C-F, Fig. S4B). To determine whether these effects depended on NPF expression, we simultaneously activated *NPF-Gal4* neurons and knocked down *NPF* using *UAS-NPF-RNAi*. This manipulation restored feeding and Margin Zone huddling to dormancy-like levels, implicating NPF expression in the regulation of both behaviors during dormancy (Fig. 4C-F, Fig S4B).

### NPF^FSB^ Activation Suppresses Dormancy-Associated Margin Zone Huddling

*NPF-Gal4* labels roughly 50 neurons in the brain and a group of enteroendocrine cells in the midgut that collectively regulate action selection for diverse processes including hunger, arousal, and feeding (Fig. 4G) (36, 39, 40). As the function of NPF cells varies with anatomy, we sought to further isolate the neurons for the feeding and huddling phenotypes produced by broad NPF activation. We used TRPM8 to activate anatomically defined subsets of *NPF*-expressing cells, including NPF enteroendocrine cells in the gut, LNds, P1s, and fan-shaped body neurons (41–45). None of these subsets reproduced warm-like feeding or Feeding Zone grouping observed with broad NPF activation (Fig. 4I, Fig. S4C-D). In contrast, activation of ∼32 *NPF*-expressing neurons innervating the fan-shaped body (NPF^FSB^) significantly reduced Margin Zone huddling (Fig. 4J). Importantly, activation of both the broad NPF population and NPF^FSB^ neurons also reduced total huddle participation independent of arena region (Fig. S4E), indicating that these manipulations suppress huddling itself rather than shifting huddles toward food. Thus, NPF^FSB^ activation selectively suppresses huddling without increasing feeding, establishing that huddling propensity can be experimentally dissociated from appetitive behavior (Fig. 4K).

## Discussion

Physiological states coordinate diverse metabolic, endocrine, and behavioral processes, yet the extent to which these processes share regulatory control remains an important question. Internal states such as hunger, arousal, and reproductive motivation influence how animals process sensory information and select among competing behaviors, allowing behavioral priorities to change with physiological need and environmental conditions (7, 9, 10). Here, we identify huddling as a behavioral signature of *Drosophila* dormancy and demonstrate that this state reorganizes social behavior along two experimentally separable dimensions: the propensity to form huddles and the selection of aggregation sites. Dormancy increased huddling even in the absence of food while shifting aggregation toward the arena margins. Moreover, appetite-regulating neuropeptide manipulations dissociated feeding from huddling, demonstrating that these behavioral outputs are independently modifiable.

The identification of dormancy-associated huddling also contributes to a more comprehensive characterization of the dormant state. Reproductive arrest during cold exposure shares features with ovarian responses to other environmental stressors, suggesting that reproductive dormancy may reflect a generalized stress response rather than a specialized physiological program (6). However, reproductive arrest alone cannot distinguish a coordinated physiological state from a generalized stress response, highlighting the need for additional behavioral and physiological criteria (7). Huddling provides such a criterion because it increases despite profound locomotor suppression, does not require food, and is sensitive to olfactory signaling. Its selective suppression through neuronal activation further supports the presence of regulated behavioral organization during dormancy, rather than a collection of passive responses to cold stress.

Our findings further indicate that dormancy alters feeding homeostasis beyond the reduction in food consumption expected from lower metabolic demand. Dormant flies were less likely to initiate feeding but did not compensate by consuming larger meals, demonstrating that reduced consumption reflects suppression of meal initiation rather than reduced consummatory capacity. Upon recovery to warm conditions, flies exhibited both hyperphagia and reduced starvation resistance despite continuous access to food during dormancy, suggesting a mismatch between energetic state and appetite. Reduced starvation resistance upon recovery could reflect insufficient energy reserves, an increased energetic cost associated with reactivation of warm-temperature physiology, altered accessibility or mobilization of stored nutrients, or some combination of these mechanisms. Previous reports of elevated lipid and carbohydrate stores during *Drosophila* dormancy make the latter possibilities particularly interesting, but direct measurements of energy stores and metabolic expenditure during recovery will be required to distinguish among them.

Dormancy fundamentally alters the relationship between energetic need and appetitive motivation. Dormancy-associated appetite suppression was not equivalent to ordinary satiety.

Across multiple experiments, dormant flies consistently exhibited substantially greater Margin Zone huddling than any of the hunger or satiety states induced in warm flies. Neither prolonged starvation nor refeeding after starvation reproduced this behavior, even when recently refed flies were selected for substantial meal consumption. Nevertheless, appetitive state clearly influenced aggregation site: starved warm flies positioned themselves closer to food, and labellectomy progressively reduced Margin Zone huddling during dormancy despite leaving the food source and external environment unchanged. Thus, aggregation site selection remains sensitive to appetitive state during dormancy, but the relationship is shifted beyond the range produced by normal fluctuations in hunger and satiety. Dormancy-associated appetite suppression therefore appears to represent a distinct homeostatic condition beyond the satiated extreme of normal feeding homeostat.

The distinction between huddling propensity and aggregation site selection has implications for the organization of social behavior more generally. Aggregation is widespread among animals and can serve diverse functions, including resource exploitation, predator avoidance, and protection from environmental stress (46, 47). Although decisions about whether to associate with conspecifics and where to aggregate are often correlated, particularly around shared resources, our experiments demonstrate that these dimensions can be experimentally separated. Dormancy increased the propensity to huddle even when food was absent, whereas appetitive manipulations altered the distribution of huddles relative to food. Social aggregation may therefore emerge from separable influences on group formation and spatial preference, allowing animals to maintain social associations while adjusting resource seeking as physiological conditions change.

Targeting distinct appetite-regulating neuropeptide pathways revealed that feeding and huddling can be differentially regulated during dormancy. Broad activation of NPF neurons increased feeding while suppressing Margin Zone huddling, whereas SIFamide activation increased feeding without significantly altering huddling. Conversely, other manipulations reduced Margin Zone huddling without detectable increases in feeding. Changes in feeding did not consistently predict changes in huddling. Anatomical subdivision of the NPF population further revealed that activation of fan-shaped body NPF neurons selectively suppressed huddling without restoring warm-like feeding, consistent with functional specialization among NPF populations (36). Although these experiments establish the sufficiency of neuronal activation, they do not demonstrate that these neurons are necessary for huddling or that their endogenous activity changes during dormancy. Identifying their physiological responses and downstream targets will be necessary to establish their role in dormancy regulation. This functional dissociation also extended to reproductive physiology, the defining feature traditionally used to characterize *Drosophila* diapause. Several appetite-regulating neuropeptide manipulations increased mature egg production during dormancy without corresponding changes in feeding or huddling. Thus, even reproductive arrest can be experimentally modified without necessarily restoring other dormancy-associated phenotypes, supporting a model in which dormancy encompasses partially separable physiological and behavioral outputs.

The possibility that these outputs remain coordinated through shared state-regulating circuitry is particularly intriguing in the context of the circadian system. In addition to the profound suppression of locomotion, we observed an advance of the evening activity peak during dormancy (Fig. 1A), consistent with recent observations of altered circadian organization during *Drosophila* diapause (8). The timing of evening activity is actively set by E cells, including the LNds and fifth s-LNv (48, 49), with the E2 subclass contributing to evening anticipation and phase adjustment (50, 51) . Environmental regulation of these neurons provides a potential connection to dormancy: photoperiodic adjustments involve PDF signaling to NPF-expressing clock neurons (50, 52), while ablation of NPF-positive clock neurons advances and attenuates the evening peak (53). Cold also advances evening activity through molecular clock regulation and can recruit LNd-derived NPF signaling to reshape evening locomotion (54, 55). Together with our previous finding that E-cell activation stimulates egg production during reproductive dormancy (28), these observations raise the possibility that E-cell circuitry contributes to the coordination of circadian, reproductive, and social components of dormancy. Although activation of NPF-expressing LNds alone was insufficient to suppress huddling, this does not exclude a broader role for E-cell populations or distributed signaling among circadian and neuroendocrine circuits.

These findings also raise questions about the adaptive significance of dormancy-associated huddling. Aggregation during adverse environmental conditions is widespread among animals and can provide benefits ranging from microclimate modification to reduced energetic expenditure and water loss. Whether *Drosophila* huddling provides similar benefits remains unknown. By establishing huddling as a behavioral hallmark of dormancy and separating its regulation from aggregation site selection, our findings provide a framework for understanding how a physiological state is translated into coordinated but independently controllable behavioral outputs.

## Materials and Methods

### Fly Stocks and Husbandry

The following Gal4 driver lines were obtained from the Bloomington Drosophila Stock Center (BDSC): *ort^-/-^, ninaE^-/-^* (#1946), *elav-Gal80* (#607940), *CCHa2-Gal4* (#84602), *Lk-Gal4* (#51993), *MIP-Gal4* (#51984), *AstA-Gal4* (#51979), *Hugin-Gal4* (#58769), *sNPF-Gal4* (#51991), *AstC-Gal4* (#52017), *NPF-Gal4* (#25681), *Crz-Gal4* (#51976), *SIFa-Gal4* (#84690), and *ILP2-Gal4* (#37516).

*UAS-TNT* (#28997) and *UAS-NPF-RNAi* (#27237) were also obtained from BDSC. The NPF^FSB^ population was targeted using split-Gal4 line SS63089 (BDSC #86977), NPF^P1^ neurons using SS01587 (BDSC #75917), and NPF^Gut^ cells using *NPF-Gal4* combined with *elav-Gal80*. NPF^LNd^ (*DAL2-Gal4*) was provided by Dr. Ann-Shyn Chiang (National Tsing Hua University, Taiwan), *Orco*^-/-^ were provided by Dr. Craig Montell (University of California-Santa Barbara, USA), *UAS-TRPM8* was provided by Dr. Benjamin White (National Institutes of Health, USA). Isogenic *w1118* flies were used to generate genetic controls.

Flies were reared on standard cornmeal-agar medium at 25°C and 65% relative humidity under a 12:12 h light:dark cycle. Virgin females were collected within 2 hr of eclosion and maintained at 25°C or transferred immediately to dormancy-inducing 12°C under otherwise identical conditions. On day 3 post-eclosion, groups of 10 flies were transferred to behavioral arenas and acclimated for approximately 18 hr before recording on day 4. All behavioral assays used groups of 10 females unless otherwise indicated.

### Behavioral Recording and Tracking

Behavioral recordings were performed using the Long-Term Observation Overhead Camera (LOOC) platform (11). Custom clear acrylic arenas (6.5 cm inner diameter × 0.25 cm height) had gas-permeable Mylar membrane floors (MP Biomedicals, SKU 097640205) and a central cornmeal-agar food disc (1 cm diameter × 0.25 cm thick) unless otherwise indicated. Arenas were maintained at 12°C or 25°C, 65% relative humidity, and a 12:12 h light:dark cycle. Continuous uncompressed infrared videos were acquired at 800 × 800 pixels and 10 frames/s using Raspberry Pi cameras. Unless otherwise indicated, analyses used the 24-hr interval following acclimation.

Videos were analyzed using T-Rex (13) for individual tracking and locomotor velocity. Position data were processed using custom LOOCAPT software to quantify inter-fly distance, grouping, huddle location, and regional occupancy. Feeding was independently quantified using SLEAP (19).

### LOOCAPT Analysis and Huddle Identification

LOOC videos were sampled at one frame per minute for 24-hr recordings or one frame per 10 s for shorter recordings. Backgrounds were removed in FIJI (56) by subtracting the average frame, and the resulting images were binarized in LOOCAPT. Objects were characterized by centroid coordinates, area, perimeter, and major and minor axis lengths. A k-nearest-neighbor classifier trained on approximately 15,000 manually annotated objects identified objects containing zero, one, or two flies; rare objects containing three or more flies were manually annotated.

Pairwise distances between objects were calculated, retaining distances below 1 cm. Objects separated by less than 1 cm were assigned to groups, and overlapping groups were iteratively merged over five iterations. Group membership was defined by connected objects, such that each member was within 1 cm of at least one other member. Groups contained at least three flies, with members connected through inter-fly distances below 1 cm (∼2.5 body lengths). Group centroids were calculated from the mean coordinates of their members.

To establish the minimum huddle size, 10,000 randomized frames were generated by sampling 10 fly positions with replacement from normalized coordinates of warm-reared Canton-S flies. These were resampled into 10,000 simulated trials matching the duration of experimental recordings. Group frequencies were compared between observed and randomized trials across group sizes. Five flies was the smallest group size for which the observed frequency exceeded the 95th percentile of the randomized distribution and was therefore adopted as the minimum huddle size (Fig. 1J).

Coordinates were normalized to the center of the food disc, or arena center in food-free trials. Group centroids were assigned to 1-cm spatial bins. The food disc had a radius of 0.5 cm. The Feeding Zone was defined as an annulus with an interior radius of 0.5 cm and an exterior radius of 1.15 cm centered on the food disc. The exterior radius was defined as the food disc radius plus approximately two fly body lengths. The remaining arena area constituted the Margin Zone; an annulus with interior radius of 1.15 cm and an exterior radius of 3.25 cm. For food-proximity analyses (Fig. S4B,C), the distance of each fly from the edge of the central food disc was calculated at each sampled time point, irrespective of grouping behavior. Distances were averaged across all 10 flies and the 24-hr recording period to obtain one mean distance per independent trial.

### Caffeine Treatment and Sleep Deprivation

For caffeine treatment, caffeine powder (MilliporeSigma, C0750) was incorporated into standard cornmeal-agar food at a final concentration of 5 mg/mL. Dormant flies were exposed to caffeine beginning upon transfer to behavioral arenas approximately 18 hr before recording. Control flies received standard food without caffeine. Groups of 10 flies were recorded continuously for 24 hr at 12°C.

For sleep deprivation, warm-reared flies were subjected to 12 hr of mechanical agitation using a custom-built agitator under constant illumination from 10 PM to 10 AM, corresponding to the expected dark phase. Control flies were maintained in vials within a dark enclosure in the same room without agitation. Immediately following treatment, groups of 10 flies were transferred to behavioral arenas and recorded for 60 min at 25°C.

### SLEAP Feeding Analysis

SLEAP v1.6.5 was trained using images sampled at 1 frame/s from warm and dormant Canton-S recordings. Approximately 22,500 instances of flies were annotated at two landmarks: the center of the head (Head) and anterior third of the abdomen (Body). Feeding was classified when the Head landmark was within 0.13 cm of the current food edge. For manual validation, feeding was scored when the proboscis or most of the head contacted the food. Four hours of manually annotated video were compared with SLEAP predictions using paired Student’s *t* tests.

### Expresso Assay

Feeding was quantified using Expresso (21). Individual flies were deprived of food for 24 hr at 25°C, 12°C, or following transfer from 12°C to 25°C (recovery), with water supplied using saturated Kimwipes. At approximately ZT8, flies received 300 mM sucrose for 30 min. Meal initiation and volume were quantified using Expresso acquisition and analysis software.

### Starvation Resistance

Groups of 3-5 flies were maintained in food-free vials containing water-saturated Kimwipes at their respective temperatures (12°C or 25°C), 65% relative humidity, and a 12:12 h light cycle. Recovery flies were transferred directly from 12°C to 25°C at starvation onset. Survival was scored every 6 hr, and surviving flies were transferred to fresh vials following each death to limit cannibalism and microbial growth.

### Satiety Manipulations

Warm-reared flies were continuously fed (Fed), deprived of food for 24 hr (Starved), or deprived of food for 24 hr followed by 10 min access to 100 mM sucrose containing Royal Blue Food Color (AmeriColor, Placentia, CA, USA) (Refed). Food deprivation was performed in groups of 20 with water-saturated Kimwipes. Refed flies were selected for abdominal distension indicating recent feeding. Groups of 10 flies were immediately transferred to behavioral arenas and recorded at 25°C.

### Labellectomy

Flies were anesthetized with CO₂ followed by ice, and the labellum was removed by transecting the distal proboscis at the hyoid sclerite of the prepharynx using Vannas spring scissors (Fine Science Tools). Surgeries were completed within 20 min and verified visually. Sham-operated flies underwent identical anesthesia and handling without transection. Groups of 10 flies recovered directly in behavioral arenas at 12°C and were recorded continuously for 72 hr. Trials with any mortality during recording were excluded.

### Neuropeptide Manipulations

Five putatively anorexigenic neuronal populations (CCHa2, Lk, MIP, AstA, and Hugin) were inhibited using *UAS-TNT*, whereas six putatively orexigenic populations (sNPF, AstC, NPF, Crz, SIFa, and ILP2) were activated using *UAS-TRPM8*. Corresponding Gal4/+ and UAS/+ controls were generated using isogenic *w1118*. Experimental and control females were transferred to 12°C at eclosion and recorded for 24 hr from ZT0 on day 4 to ZT0 on day 5. Huddling and feeding were quantified using LOOCAPT and SLEAP, respectively.

For reproductive measurements, females were collected immediately following recording at ZT0 on day 5. Ovaries from 20-40 females per genotype were dissected, and mature stage 14 eggs were counted separately for each ovary.

For simultaneous activation and NPF knockdown, *UAS-NPF-RNAi* and *UAS-TRPM8* were coexpressed using *NPF-Gal4*. Gut, P1, LNd, and fan-shaped body-associated NPF populations were activated using their corresponding Gal4 or split-Gal4 drivers with *UAS-TRPM8* under identical dormancy-inducing conditions.

### Immunohistochemistry and Confocal Microscopy

Adult brains, ventral nerve cords, and gastrointestinal tracts were dissected and immunostained as previously described (Cui et al., 2024). Briefly, tissues were dissected in phosphate-buffered saline (PBS), fixed in 4% paraformaldehyde (Electron Microscopy Sciences, #15711) for 15-25 min, washed in 0.1% PBS-Triton X-100 (PBT), and blocked in 5% normal goat serum. Samples were incubated with primary antibodies for 2-5 days at 4°C, followed by secondary antibodies for 24-48 hr at 4°C. Primary antibodies included chicken anti-GFP (1:3,000; Abcam, #ab13970) and mouse anti-Bruchpilot (nc82; 1:20; Developmental Studies Hybridoma Bank, #Nc82). Secondary antibodies included Alexa Fluor 488-conjugated goat anti-chicken IgG (1:1,000; Invitrogen, #A11039) and Cy5-conjugated goat anti-mouse IgG (1:500; Invitrogen, #A10524). Gastrointestinal tissues were additionally stained with phalloidin (Sigma-Aldrich, #P2141) to visualize F-actin. Samples were mounted in SlowFade medium (Thermo Fisher Scientific, #S36936) and imaged using a Leica STELLARIS confocal microscope at the University of Nevada, Las Vegas Imaging Core.

### Statistical Analysis

Statistical analyses were performed using GraphPad Prism 11.1.0 (GraphPad Software, Boston, MA, USA). Normality was assessed using Shapiro-Wilk tests for *n* < 50 and Kolmogorov-Smirnov tests for *n* ≥ 50. Variance equality was assessed using F tests for two groups and Brown-Forsythe tests for three or more groups. Normally distributed data with equal variances were analyzed using unpaired Student’s *t* tests or one- or two-way ANOVA, as appropriate. Welch’s *t* tests were used for unequal variances, and Mann-Whitney U or Kruskal-Wallis tests with Dunn’s correction were used for nonnormal data. One-way ANOVAs were followed by Dunnett’s test for comparisons against designated controls or Tukey’s test for pairwise comparisons among groups, as indicated in the figure legends. Survival was analyzed using Kaplan-Meier curves and Mantel-Cox log-rank tests. Statistical significance was set at *P* < 0.05, with multiple-comparison adjustments as indicated. Prespecified pairwise Mann-Whitney U tests were adjusted for multiple comparisons using the Holm-Šídák procedure. Data are presented as mean ± SEM unless otherwise noted.

## Acknowledgments

We thank Kiara Concepcion, Kamonkan (Joy) Gamnerdsiri, Nelly Spartalyan, and Kim Nguyen for assistance with video processing and training the SLEAP feeding classifier.

## Author Contributions

S.R.O. and M.R.M. designed research; S.R.O., R.J., J.L.M., J.D.-R., and M.R.M. performed research; N.Y. contributed resources; S.R.O. and M.R.M. analyzed data; and S.R.O. and M.R.M. wrote the paper.

## Data Availability

Data supporting the findings of this study and custom analysis code will be made publicly available upon publication. Repository information and accession links will be provided prior to publication.

## Supporting Information

**Fig. S1.**
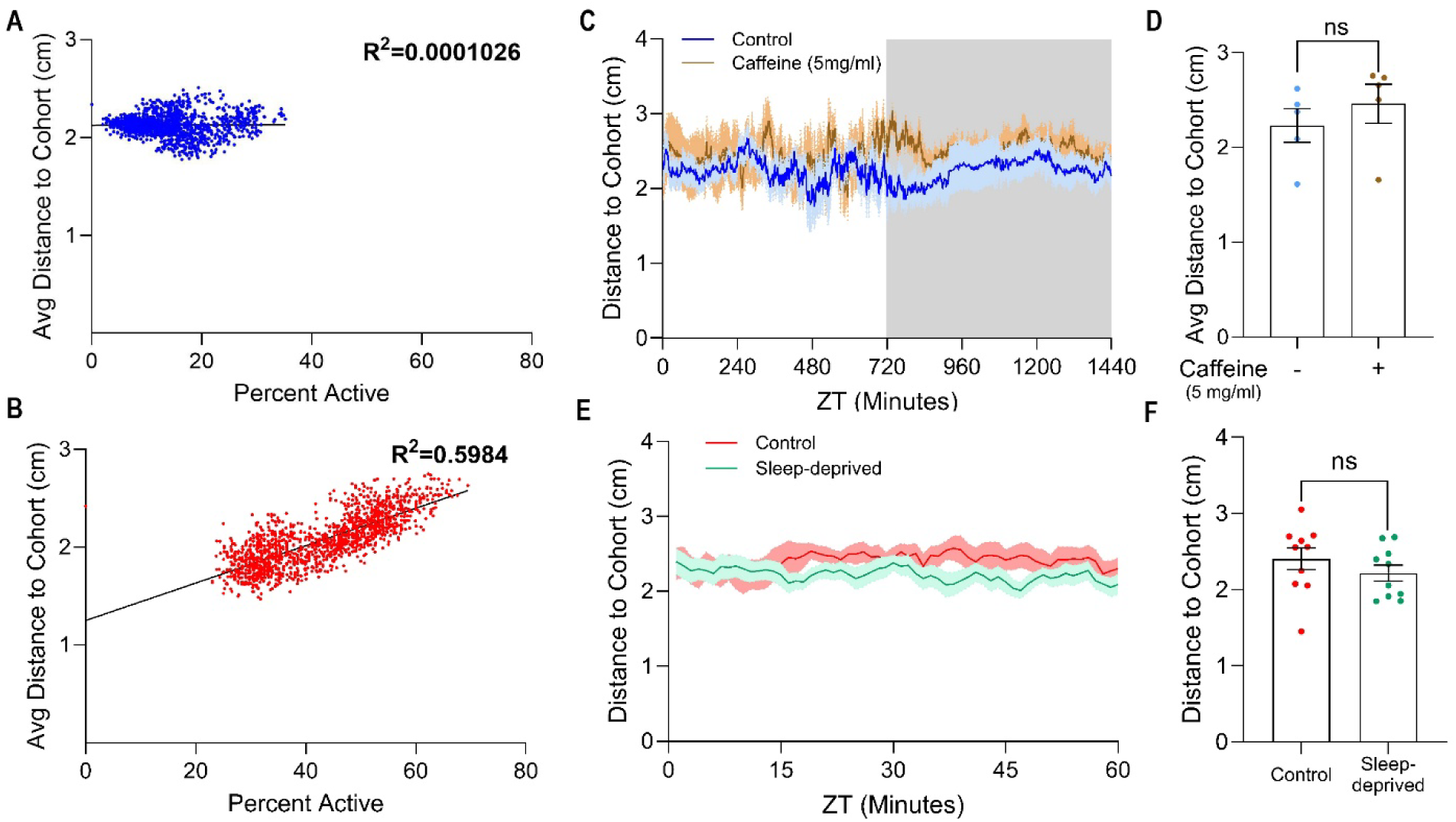
Dormancy disrupts the relationship between locomotor activity and social spacing. **(A and B)** Relationship between locomotor activity and mean distance to cohort at dormancy-inducing 12°C (**A**, blue) or 25°C (**B**, red). Each point represents one 1-min interval pooled across 15 independent trials (21,600 intervals per temperature). Lines indicate linear regression; R² values are shown on each graph. **(C)** Mean distance to cohort of dormant flies maintained at 12°C with (brown) or without (blue) 5 mg/ml caffeine over a 24-hr 12:12 light:dark cycle. Lines indicate mean and shading indicates ± SEM (*n* = 5 trials per condition). Gray shading indicates the dark phase. **(D)** Mean distance to cohort over the 24-hr recording period with or without 5 mg/ml caffeine. Each point represents one independent trial (*n* = 5 trials per condition); bars indicate mean ± SEM. Student’s *t* test, *ns*. **(E)** Mean distance to cohort of warm-reared control (red) or sleep-deprived (green) flies during the 60-min assay. Lines indicate mean and shading indicates ± SEM (*n* = 10 trials per condition). **(F)** Mean distance to cohort over the assay. Each point represents one independent trial (*n* = 10 trials per condition); bars indicate mean ± SEM. Student’s *t* test, *ns*.

**Fig. S2.**
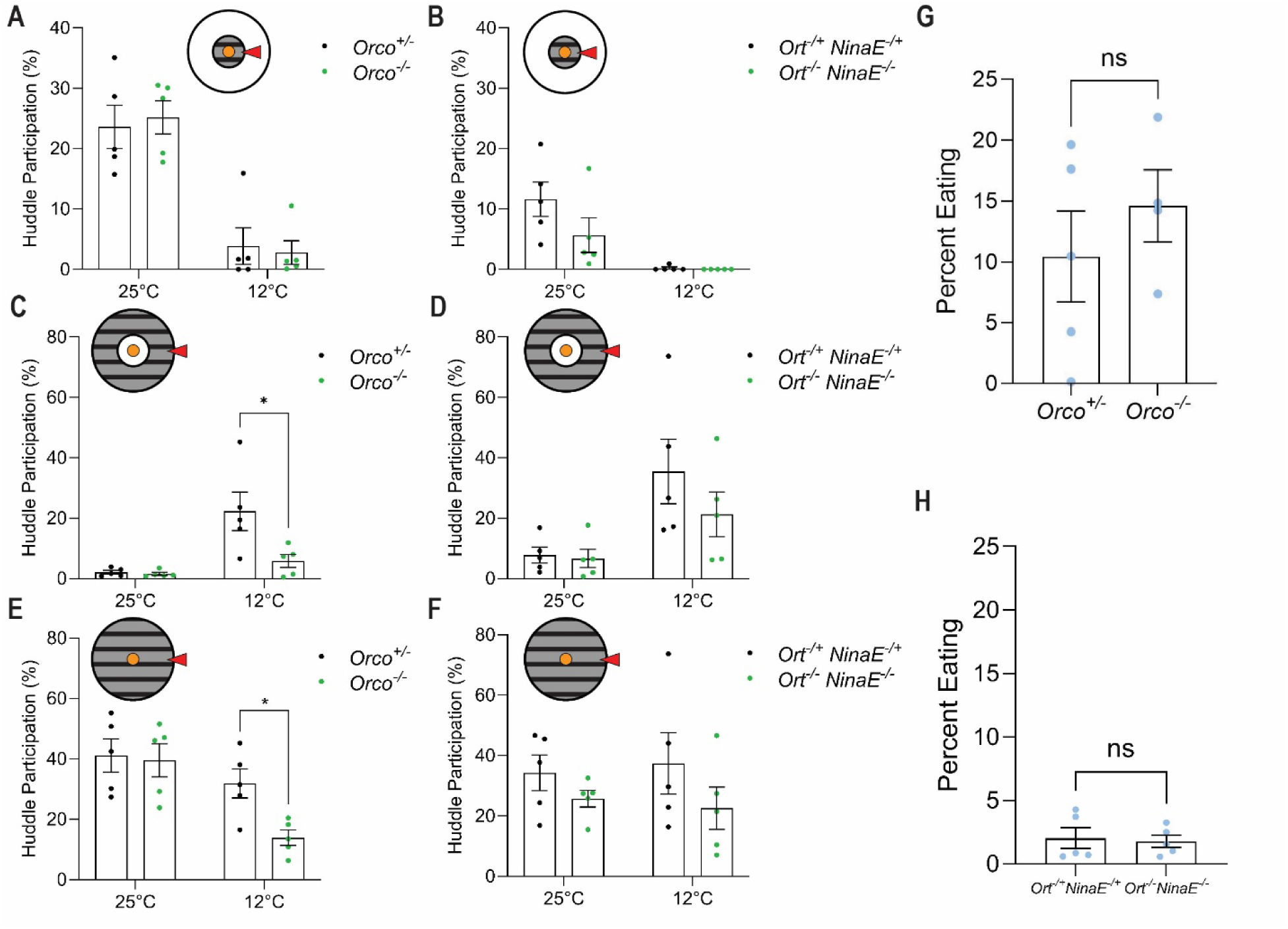
Olfactory, but not visual, signaling contributes to dormancy-associated huddling. **(A, C, E)** Huddle participation in *Orco* heterozygous control (*Orco*^+/−^) and anosmic *Orco*^−/−^ flies maintained at 25°C or dormancy-inducing 12°C at locations indicated by red arrows and shaded insets. **(A)** Feeding Zone huddle participation. **(C)** Margin Zone huddle participation. **(E)** Total huddle participation independent of arena region. **(B, D, and F)** Huddle participation in visually intact control (*ort*−/+; *ninaE*−/+) and visually impaired *ort*^−/−^; *ninaE*^−/−^ flies maintained at 25°C or 12°C at locations indicated by red arrows and shaded insets. **(B)** Feeding Zone huddle participation. **(D)** Margin Zone huddle participation. **(F)** Total huddle participation independent of arena region. Each point represents one independent trial (*n* = 5 trials per genotype and temperature); bars indicate mean ± SEM. Student’s *t* tests comparing mutant and corresponding control genotypes at each temperature; *, *P* < 0.05. **(G and H)** Mean percentage of time spent feeding over 24 hr in *Orco* heterozygous controls and anosmic mutants **(G)**, or visually intact controls and visually impaired *ort; ninaE* mutants **(H)**, maintained at 12°C and quantified using SLEAP. Each point represents one independent trial (*n* = 4-5 trials per genotype); bars indicate mean ± SEM. Student’s *t* test, *ns*.

**Fig. S3.**
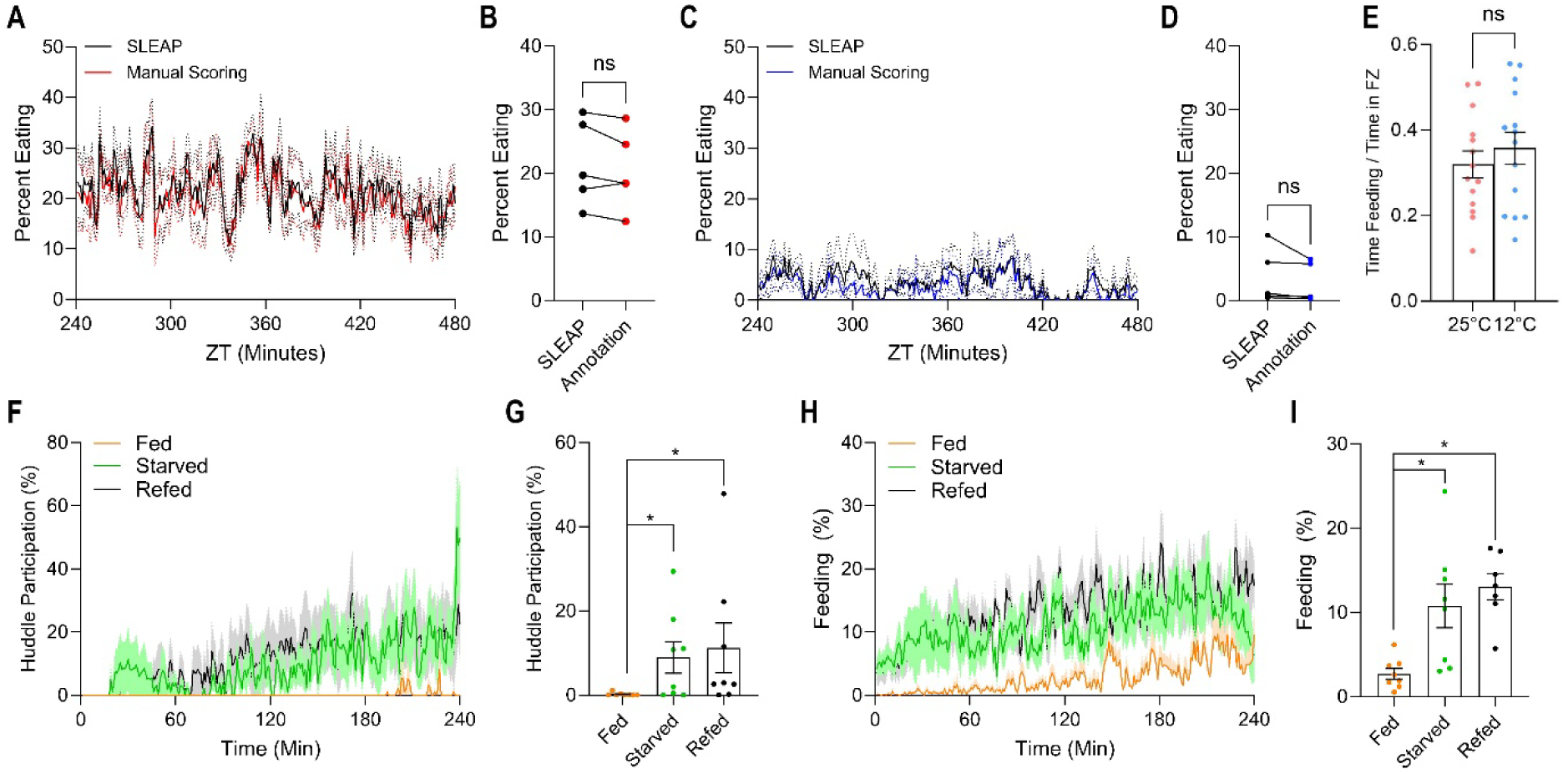
Appetitive state regulates feeding and food-associated huddling. **(A and C)** Comparison of automated SLEAP-based feeding quantification (black) with blinded manual scoring of feeding at 25°C (**A**, red) or dormancy-inducing 12°C (**C**, blue) over a 4-hr interval. Lines indicate mean and dashed lines indicate ± SEM. **(B and D)** Mean percentage of flies feeding as quantified by SLEAP or manual annotation at 25°C **(B)** or 12°C **(D)**. Connected points represent paired measurements from the same independent trial. Paired Student’s *t* test, *ns*. **(E)** Proportion of time spent feeding relative to total time spent in the Feeding Zone, quantified using SLEAP. Each point represents one independent trial (*n* = 15 trials per temperature); bars indicate mean ± SEM. Student’s *t* test, *ns*. **(F-I)** Effect of acute changes in satiety on Feeding Zone huddling and feeding in warm-reared flies. Flies were continuously fed (Fed), food deprived for 24 hr (Starved), or food deprived for 24 hr and allowed to feed for 10 min on 100 mM sucrose containing blue food coloring (Refed). For the Refed condition, flies exhibiting the greatest abdominal distension and visible blue coloration were selected to enrich for individuals that had recently consumed a large meal. **(F)** Percentage of flies participating in Feeding Zone huddles during the 4-hr assay. Lines indicate mean and shading indicates ± SEM. **(G)** Mean Feeding Zone huddle participation over the assay. Each point represents one independent trial; bars indicate mean ± SEM. Mann-Whitney U tests comparing Fed with Starved and Refed, with Holm-Šídák correction for two comparisons; *, *P* < 0.05. **(H)** Percentage of flies feeding during the 4-hr assay, quantified using SLEAP. Lines indicate mean and shading indicates ± SEM. **(I)** Mean percentage of flies feeding over the assay. Each point represents one independent trial; bars indicate mean ± SEM. Mann-Whitney U tests with Holm-Šídák correction for two comparisons, *, *P* < 0.05.

**Fig. S4.**
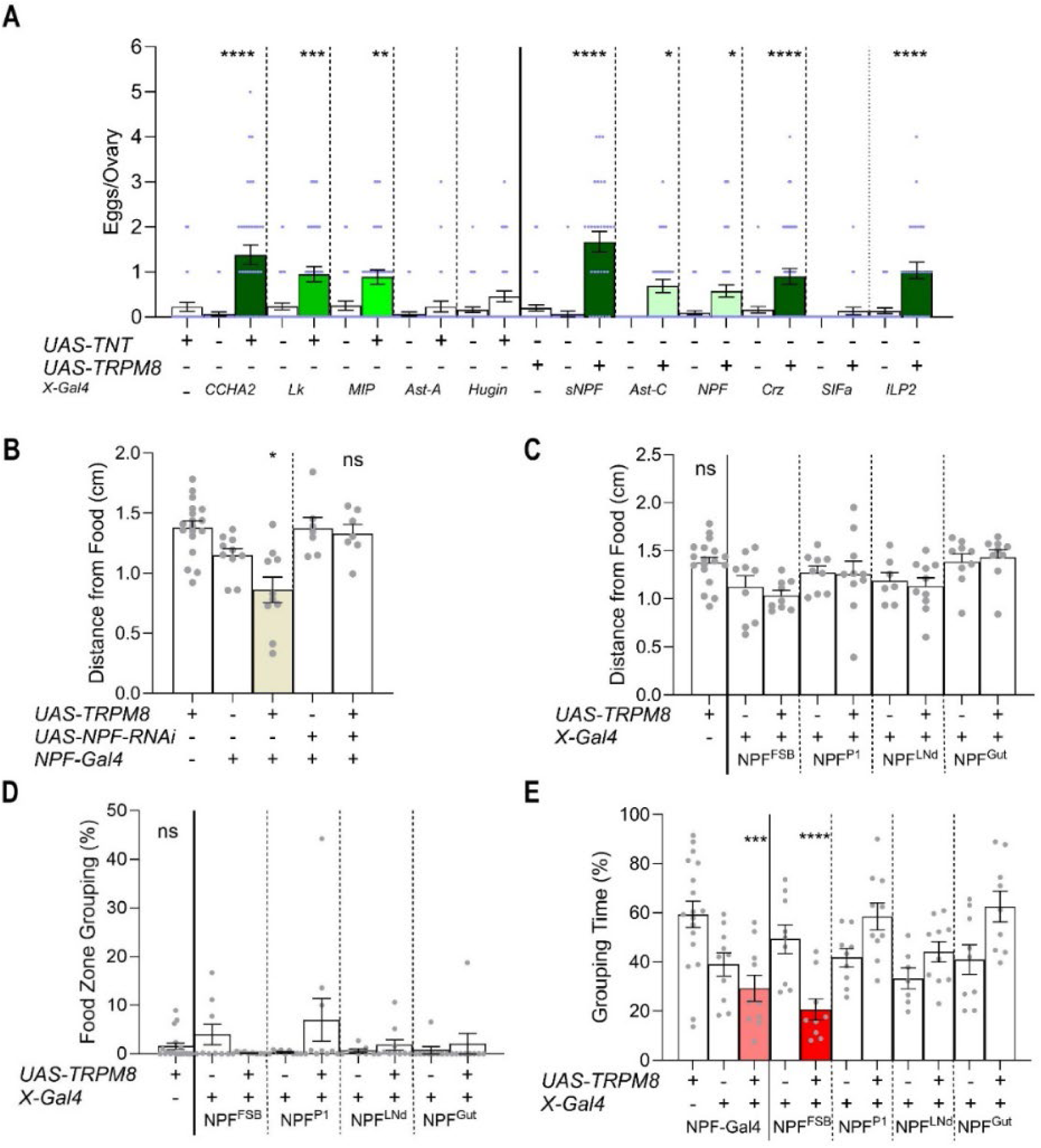
Appetite-regulating neuropeptide pathways differentially affect reproductive and behavioral outputs of dormancy. **(A)** Mature egg production in flies maintained at dormancy-inducing 12°C following inhibition of putatively anorexigenic pathways (CCHa2, Lk, MIP, AstA, and Hugin) using *UAS-TNT*, or activation of putatively orexigenic pathways (sNPF, AstC, NPF, Crz, SIFa, and ILP2) using *UAS-TRPM8*. Each point represents one ovary; bars indicate mean ± SEM. One-way ANOVA followed by Dunnett’s multiple-comparisons test against the corresponding UAS controls. Bar fill indicates significance: white, *ns*; pale green, *P* < 0.05; bright green, *P* < 0.01; green, *P* < 0.001; dark green, *P* < 0.0001. **(B and C)** Effects of broad NPF activation and simultaneous NPF knockdown (**B**), or activation of anatomically defined NPF subsets (**C**), on proximity to food during dormancy. Mean distance from the food edge for all flies over the 24-hr recording period, irrespective of grouping behavior. Each point represents one independent trial; bars indicate mean ± SEM. **(B)** One-way ANOVA followed by Tukey’s multiple-comparisons test. Symbols indicate significance relative to both corresponding genetic controls, with the less significant adjusted comparison reported: *, *P* < 0.05; **, *P* < 0.01; ***, *P* < 0.001; ****, *P* < 0.0001; ns. **(C)** One-way ANOVA followed by Dunnett’s multiple-comparisons test against the corresponding UAS controls; *ns*; ****, *P* < 0.0001. **(E)** Mean Feeding Zone huddle participation over 24 hr following activation of anatomically defined NPF subsets (NPF^FSB^, NPF^P1^, NPF^LNd^, and NPF^Gut^) using *UAS-TRPM8*. Huddles were defined as groups of five or more flies with centroids located within the Feeding Zone. Each point represents one independent trial; bars indicate mean ± SEM. One-way ANOVA, *ns*. **(F)** Mean total huddle participation over 24 hr, independent of arena region, following activation of broad NPF neurons or the indicated NPF subsets using *UAS-TRPM8*. Huddles were defined as groups of five or more flies regardless of location. Each point represents one independent trial; bars indicate mean ± SEM. One-way ANOVA followed by Dunnett’s multiple-comparisons test against the corresponding UAS controls. Bar fill indicates significance: white, *ns*; light red, *P* < 0.001; dark red, *P* < 0.0001.

